# Average oxidation state of carbon in proteins

**DOI:** 10.1101/004127

**Authors:** Jeffrey M. Dick

## Abstract

The degree of oxidation of carbon atoms in organic molecules depends on the covalent structure. In proteins, the average oxidation state of carbon (*Z*_C_) can be calculated as an elemental ratio from the chemical formula. To investigate oxidation-reduction (redox) patterns, groups of proteins from different subcellular locations and phylogenetic divisions were selected for comparison. Extracellular proteins of yeast have a relatively high oxidation state of carbon, corresponding with oxidizing conditions outside of the cell. However, an inverse relationship between *Z*_C_ and redox potential occurs between the endoplasmic reticulum and cytoplasm; this trend is interpreted as resulting from overall coupling of protein turnover to the formation of a lower glutathione redox potential in the cytoplasm. In Rubisco homologues, lower *Z*_C_ tends to occur in organisms with higher optimal growth temperature, and there are broad changes in *Z*_C_ in whole-genome protein compositions in microbes from different environments. Energetic costs calculated from thermodynamic models suggest that thermophilic organisms exhibit molecular adaptation to not only high temperature but also the reducing nature of many hydrothermal fluids. A view of protein metabolism that depends on the chemical conditions of cells and environments raises new questions linking biochemical processes to changes on evolutionary timescales.

## 1 Introduction

Chemical reactions involving the transfer of electrons, known as oxidation-reduction or redox reactions, are ubiquitous in cellular and environmental systems [1, 2]. In the cell, the oxidation of thiol groups in proteins to form disulfides has the potential to regulate (activate or inhibit) enzymatic function [3]. Because these reactions are reversible on short timescales, a regulatory network known as redox signalling is made possible by reactions of small-molecule metabolites, including glutathione (GSH) and reactive oxygen species [4]. On timescales of metabolism, complex oxidation-reduction reactions are required for the formation (anabolism) and degradation (catabolism) of proteins and other biomolecules. Although many individual steps in biomass synthesis are irreversible, much biomass is ultimately recycled through endogenous metabolism [5]. On longer timescales, forces outside of individual cells and organisms sustain the redox disequilibria between inorganic and/or organic species that provide the energy source for metabolisms suited to a multitude of environments [6]. In turn, the actions of organisms can alter the redox conditions on Earth; the oxygenation of the atmosphere and oceans over geological time has a biogenic origin, and changed the course of later biological evolution [7].

Through evolution, the sequences of genes, and their protein products, are progressively altered. The elemental stoichiometry (chemical formula) and standard Gibbs energy of the molecules have a primary impact on metabolic requirements for energy and elemental resources. The energetic cost for synthesis of biomass is a function not only of the composition of the biomass, but also of environmental parameters including temperature and the concentrations of metabolic precursors. Temperature and oxidation-reduction potential have profound effects on the relative energetic costs of formation of different amino acids [8] or proteins [9]. These energetic costs are sensitive to differences in the elemental compositions of biomolecules. To a first approximation, a shift to a more reducing environment alters the energetics of reactions in a direction that favours the formation of relatively reduced chemical compounds. In a field test of this principle, metagenomic sequences for the most highly reduced proteins were found in the hottest and most reducing zones of a hot spring [10, 11].

The purpose of this study is to investigate a particular stoichiometric quantity, the average oxidation state of carbon (*Z*_C_, defined below), as a comparative tool for identifying compositional patterns at different levels of biological organization. By comparing a quantity derived from the elemental compositions of proteins, this study addresses one aspect of biochemical evolution. However, the questions raised here differ in important respects from conventionally defined biochemistry and evolutionary biology. Biochemical studies are most often concerned with the functions of molecules [12], including enzymatic catalysis and non-covalent interactions involved in the structural conformation of proteins and binding of ligands. Studies in molecular evolution often place emphasis on the historical relationships between sequences, but not their physical properties [12]. Combining these viewpoints, most current work assumes that structural stability of proteins is the primary criterion for molecular adaptation to high temperature [13]. In contrast, in this study, more attention is given to the stoichiometric and energetic demands of the reactions leading to protein formation. Because material replacement of proteins depends on metabolic outputs [14], the compositional differences among proteins have significant consequences for cellular organization and metabolism.

The following questions have been identified: 1) How does the relationship between Z_C_ of amino acids and corresponding codons relate to the origin or form of the genetic code? 2) How do the differences in *Z*_C_ between membrane proteins and others compare with properties of amino acids, e.g. hydrophobicity, known to favour localization to membranes? 3) How are the differences in *Z*_C_ of proteins among eukaryotic subcellular compartments related to differences in redox potential? 4) How are the differences in *Z*_C_ in families of redox-active proteins related to the standard reduction potentials of the proteins? 5) How are the differences in *Z*_C_ among different organisms, both in terms of bulk (genome-derived) protein composition and for homologues of a single family (Rubisco), related to environmental conditions, especially temperature and redox potential?

In the Results each problem is briefly introduced, the empirical distribution of *Z*_C_ is described, and a discussion is developed to explore how the patterns reflect biochemical and evolutionary constraints. These short discussions, corresponding to topics (1)-(4), should be regarded as preliminary, and probably incomplete interpretations. The discussions are limited because there is no complete conceptual framework that links the biochemical reactions with the evolutionary processes that are implicit in all of the comparisons. The final section of the Results goes into more detail for the phylogenetic comparisons (topic 5) by examining the relative Gibbs energies of formation of proteins in environments of differing redox potential.

## 2 Methods

Throughout this study, “reducing” and “oxidizing” are used in reference to oxidation-reduction potential, tied to a particular redox couple or to environmental conditions, often expressed as millivolts on the Eh scale. “Reduced” and “oxidized” are used to refer to variations in the oxidation state of carbon.

The formal oxidation state of a carbon atom in an organic molecule can be calculated by counting −1 for each carbon-hydrogen bond, 0 for each carbon-carbon bond, and −1 for each bond of carbon to O, N or S [8, 15]. In photosynthesizing organisms, and autotrophs in general, the carbon source is CO_2_, having the highest oxidation state of carbon (+4). The products of photosynthetic reactions include proteins and other biomolecules with a lower oxidation state of carbon. Even if the molecular structure is unknown, analytical elemental compositions can be used in calculations of the average oxidation state of carbon in biomass [16, 17]. Because any gene or protein sequence corresponds to a definite canonical (nonionized, unphosphorylated) chemical formula, the average oxidation state of carbon in these biomolecules is easily calculated.

In amino acids and proteins, the average oxidation state of carbon (*Z*_C_) can be calculated using

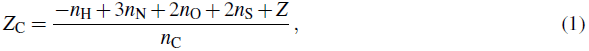

where *Z* is the charge on the molecule, and *n*_C_, *n*_H_, *n*_N_, *n*_O_ and *n*_S_ are the numbers of the subscripted elements in the chemical formula of the molecule. The coefficients on the terms in the numerator derive from formal charges of atoms other than C, as follows: H (+1), N (−3), O (−2), S (−2). Negative formal charges reflect greater electronegativities of these elements compared to carbon. If two thiol groups react to form a disulfide bond, the oxidation states of the two affected sulfur atoms change from −2 to −1. Although H_2_ is produced in this reaction, the oxidation state of carbon in the protein remains constant. It follows that equation (1) is applicable only to chemical formulas of proteins in which the N, O, and S are all fully reduced (bonded only to H and/or C).

The *Z* in equation (1) ensures that ionization by gain or loss of a proton, having an equal effect on *Z* and *n*_H_, does not change the *Z*_C_. Likewise, gain or loss of H_2_O, which affects equally the values 2*n*_H_ and *n*_O_, does not alter the average oxidation state of carbon [15]. Accordingly, the *Z*_C_ of a peptide formed by polymerization of amino acids (a dehydration reaction) is a weighted average of the *Z*_C_ in the amino acids, where the weights are the number of carbon atoms in each amino acid. As an example, the *Z*_C_ of hen egg white lysozyme, having a chemical formula of C_613_H_959_N_193_O_185_S_10_, is 0.016. This protein is oxidized compared to many other proteins, which commonly have negative values of *Z*_C_.

To aid in reproducibility, data files of protein sequences or amino acid composition, except large files available from public databases, and computer program files for the calculations are provided in the Supporting Information. The calculations and figures were generated using the R software environment [18] together with the CHNOSZ package [19].

## 3 Results

### 3.1 Comparison of *Z*_C_ of amino acids with hydropathy and properties of codons

In contemplating the ancient origin of the genetic code, the chemical similarities of respective codons and amino acids have been used to argue for coevolution (shared biosynthetic pathways) [20] or a tendency toward similar physicochemical properties. The possible advantages that were identified for similar physicochemical properties include enhancing the steric interactions between amino acids and codons [21], or increasing the similarity between different amino acids resulting from a single DNA base mutation in order to maintain protein structure [20].

In the genetic code, the first two bases (a “doublet”) are more indicative of the amino acid than the third position of the codon [21]. The *Z*_C_ of amino acids are compared with the values calculated for the corresponding RNA nucleobase doublets in figure 1*a*. Some of the doublets, e.g. UU (phenylalanine, leucine), CU (leucine), UC (serine) and CC (proline) have identical *Z*_C_ (in this case, 1.5), leading to only 5 possible values of *Z*_C_ for the doublets. The overall relationship suggested by figure 1*a* is loose correlation between the *Z*_C_ of amino acids and of the RNA doublets. The most highly reduced amino acids, leucine (L) and isoleucine (I), are coded for by doublets having the two lowest *Z*_C_ values.

**Figure 1:**
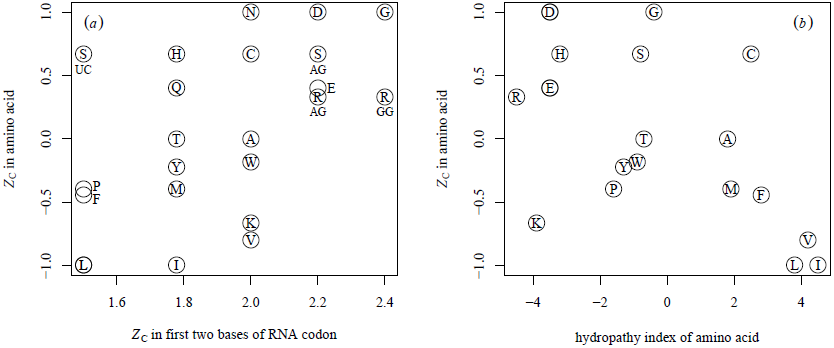
Average oxidation state of carbon (*Z*_C_) in amino acids compared with (*a*) *Z*_C_ in first two bases of the corresponding RNA codons and (*b*) hydropathy index of the amino acids taken from Kyte and Doolittle, 1982 [24]. Standard one-letter abbreviations for the amino acids are used to identify the points. In (*a*), the different codon compositions for serine (S) and arginine (R) are indicated by letters below the symbols, and some amino acid labels are shifted for readability. In (*b*), labels for asparagine (N) and glutamine (Q) are omitted for clarity; they plot at the same positions as aspartic acid (D) and glutamic acid (E), respectively.

The increase in *Z*_C_ going from leucine to alanine to glycine (figure 1) is reflected in the metastability fields of these amino acids, which occur in order of increasing oxidation potential, or oxygen fugacity [22]. Metastable equilibrium refers to the equalization of the energies of reactions to form the amino acids; it is a partial equilibrium because the amino acids generally remain unstable with respect to inorganic species. Likewise, the relative Gibbs energies of formation reactions of amino acids differ considerably between hydrothermal (hot, reducing) and surface (cool, oxidizing) environments [8]. These patterns support the possibility of metastable equilibria in hydrothermal environments between amino acids and nucleobase sequences that are paired in the genetic code. Tests of the potential for these states, carried out using Gibbs energies available at high temperature [9, 23], could reveal thermodynamic constraints on the energetics of abiotic or early biosynthesis independent of the arguments based on similarities in protein structure and biosynthetic pathways [20, 21].

**Figure 2:**
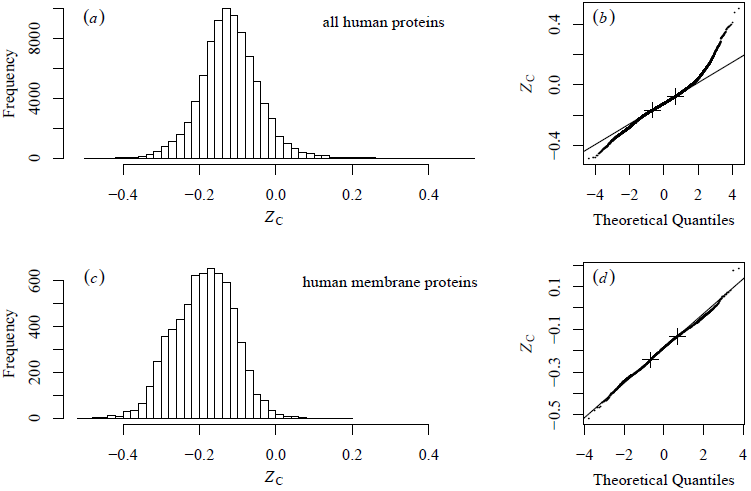
Average oxidation state of carbon shown in histograms and normal probability plots for (*a-b*) all human proteins and (*c-d*) human membrane proteins. Only proteins of sequence length greater than or equal to 50 amino acids are considered. In the normal probability plots the lines are drawn through the 1st and 3rd quartiles, indicated by the crosses.

The hydropathy index, based on the relative hydrophobicity and hydrophilicity of amino acids [24], is commonly used for identifying probable membrane-spanning domains of proteins. In figure 1*b*, *Z*_C_ is compared with the hydropathy values for individual amino acids. The three most hydropathic amino acids, isoleucine, leucine and valine, are also the three with the lowest *Z*_C_. Therefore, membrane proteins with hydrophobic domains are likely to be more reduced than other proteins. The following sections examine the actual differences in human and yeast proteins.

### 3.2 Differences in *Z*_C_ of membrane proteins

The lipid (fatty acid) components of membranes are reduced relative to many other biomolecules, including amino acids, nucleotides, and saccharides (see figure 1 of Amend *et al*., 2013 [25]). Proteins that are embedded in membranes tend to contain more hydrophobic amino acids, which enhance solubility of proteins in the membrane environment [24] and generally are relatively reduced (figure 1*b*).

To compare membrane proteins with other human proteins, sequences for all human proteins were taken from the UniProt database [26], and sequences for predicted membrane proteins were taken from all FASTA sequence files provided in Additional File 2 of Almén *et al*., 2009 [27]. Only sequences at least 50 amino acids in length were considered. The distribution of *Z*_C_ of all human proteins (figure 2*a*; *n*=83994) is centred on −0.123 (median), −0.120 (mean). In the *Z*_C_ of human membrane proteins, the distribution is shifted to lower values (figure 2*c*; *n* = 6627, −0.186 median, −0.189 mean). The mean value is lower for membrane proteins than for all human proteins (Student’s *t*-test: *p* < 2.2 × 10^‒16^). Thus, the proteins located in the membranes are, on average, more reduced than other proteins in humans. It is tempting to speculate that the coexistence of reduced proteins with other relatively reduced biomolecules (lipids) reflects a compositional similarity that would contribute to energy optimization if metabolic pathways for proteins and lipids were operating under common redox potential conditions.

The observed distributions of *Z*_C_ are each compared with a normal distribution in normal probability plots (theoretical quantile-quantile (Q-Q) plots) in figure 2*b*,*d*. The steeper trends in the low- and high-quantile range of figure 2*b* indicate that the distribution of *Z*_C_ of human proteins has relatively long tails, especially at high *Z*_C_, compared to a normal distribution. Although an asymmetry is apparent in the uneven shape of the histogram in figure 2*c* and the wiggles in figure 2*d*, the overall distribution of *Z*_C_ of the membrane proteins more closely resembles a normal distribution. Comparisons with normal distributions have implications, through the central limit theorem [28], for assessing the impact of many small-scale, independent effects in evolution on the chemical composition of organisms or their components. This theme, however, is not developed further here; instead, the overall differences in *Z*_C_ of proteins in subcellular compartments are considered next.

### 3.3 Subcellular differences in *Z*_C_ of proteins and comparison with subcellular redox potential

For some model organisms, including *Saccharomyces cerevisiae* (yeast), the identities of proteins associated with subcellular compartments are now available in databases. Here, calculations of *Z*_C_ of proteins and a comparison with independent measurements of redox potential are used to investigate the oxidation-reduction features and dynamics of cellular structure.

In a previous study, the limiting conditions for chemical transformations among proteins in subcellular compartments were quantified theoretically as a function of redox potential and hydration state [29]. In that study, the locations of proteins were taken from the the “YeastGFP” study of Huh *et al*., 2003 [30]. That dataset has the advantage that relative abundances of many of the proteins are available, but it is limited to 23 named locations in the cell. In order to consider more cellular components (including the membranes), a more extensive reference proteome is used in this study. This proteome is based on the current *Saccharomyces* Genome Database (SGD) [31] annotations combined with the Gene Ontology (GO) [32] vocabulary for the “cellular component” aspect, which describes many organelles and membranes within the cell. Major cell components were selected for comparison, and the *Z*_C_ was calculated for protein products of the genes, as summarized in table 1. The median values are also portrayed in the drawing of a yeast cell in figure 3.

**Table 1:** Summary of *Z*_C_ of proteins in subcellular locations of yeast. Numbers of proteins (*n*) in SGD associated with the indicated GO terms are listed. The numerators of the fractions denote membrane-associated proteins that are also listed as “integral to membrane” (GO:0016021); only these proteins were used in the calculations of *Z*_C_.

| cellular component | GO term | $n$ | median $Z_C$ | mean $Z_C$ |
| --- | --- | --- | --- | --- |
| cytoplasm | GO:0005737 | 2245 | -0.136 | -0.127 |
| nucleus | GO:0005634 | 2073 | -0.129 | -0.121 |
| mitochondrion | GO:0005739 | 1077 | -0.164 | -0.159 |
| endoplasmic reticulum | GO:0005783 | 435 | -0.191 | -0.192 |
| nucleolus | GO:0005730 | 263 | -0.137 | -0.128 |
| Golgi apparatus | GO:0005794 | 215 | -0.160 | -0.167 |
| cellular bud neck | GO:0005935 | 153 | -0.111 | -0.108 |
| vacuole | GO:0005773 | 175 | -0.164 | -0.163 |
| extracellular region | GO:0005576 | 95 | -0.096 | -0.098 |
| cellular bud tip | GO:0005934 | 96 | -0.113 | -0.110 |
| endoplasmic reticulum membrane | GO:0005789 | 283/338 | -0.209 | -0.211 |
| plasma membrane | GO:0005886 | 224/427 | -0.200 | -0.188 |
| mitochondrial inner membrane | GO:0005743 | 143/218 | -0.184 | -0.184 |
| vacuolar membrane | GO:0005774 | 100/145 | -0.205 | -0.196 |
| Golgi membrane | GO:0000139 | 76/121 | -0.203 | -0.200 |
| nuclear membrane | GO:0031965 | 54/67 | -0.161 | -0.139 |
| mitochondrial outer membrane | GO:0005741 | 51/92 | -0.176 | -0.177 |

**Figure 3:**
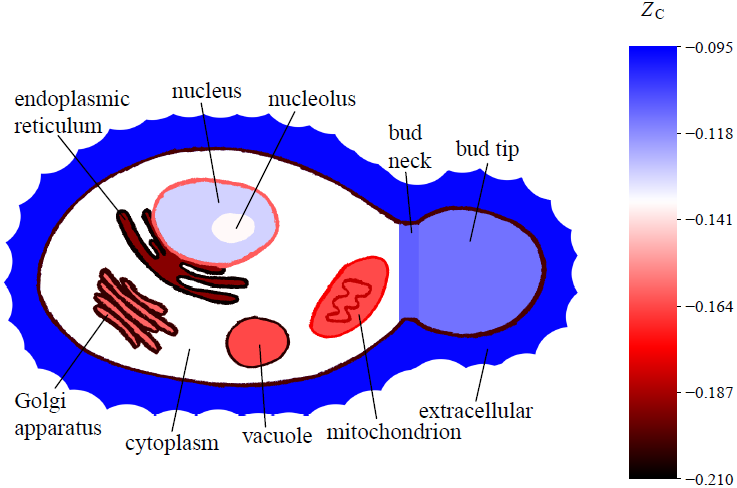
A schematic drawing of a yeast cell showing median values of the average oxidation state of carbon in proteins from selected subcellular locations listed in table 1. All cellular components listed in the table are represented in the drawing. The colour scale is adjusted so that the cytoplasm has a neutral hue (white), and locations with relatively oxidized and reduced proteins are depicted by blue and red colours, respectively. Darker red colours are used for the more reduced groups of proteins in some of the membranes such as the ER, Golgi, vacuolar and plasma membranes. The nuclear and inner and outer mitochondrial membranes are shown with lighter reds because their proteins are relatively oxidized compared to those in the other membranes. Components not separated by membranes (or with membranes not shown) include the nucleolus, bud neck and bud tip.

It is apparent that the membrane proteins are highly reduced. However, not all membranes are equal; the proteins in the nuclear and inner and outer mitochondrial membranes are less reduced than those in the plasma membrane, and the endoplasmic reticulum (ER) membrane has very highly reduced proteins. Among the organellar proteins considered, the ER has the most reduced proteins, followed by vacuoles and mitochondria; the cytoplasmic proteins are moderately reduced. The proteins in the nucleus, bud neck and bud are more oxidized than in the other compartments. The most oxidized proteins in the system are the extracellular ones. The relative *Z*_C_ of the proteins in the ER, mitochondrion, cytoplasm, and nucleus are consistent with the previous study in which the calculations took account of the abundances of proteins [29].

For comparison with *Z*_C_ of proteins, the values of subcellular redox potential (Eh, in mV) listed in table 2 were compiled from literature sources [1, 33–40]. Measurements of reduced and oxidized glutathione (GSH and GSSG) in whole-cell extracts have been interpreted as reflecting cytoplasmic redox potential, but redox-sensitive green fluorescent protein (roGFP) probes [33] provide more specific data for subcellular locations. The data are not in all cases acquired from yeast, but it has been noted that cytoplasmic Eh values based on roGFP are similar for different model organisms [36].

The redox potentials in the vacuole and extracellular space are less well constrained than other locations. Under stress response, high amounts of GSSG, but not GSH, are sequestered in vacuoles [41]. A conservative lower range for the Eh of vacuoles (−160 to −130 mV) was calculated by taking a value of 80% GSSG and computing Eh from the GSH-GSSG equilibrium at concentrations of 1–10 mM GSH (see equation 21 and figure 4 of Ref. [35]). The redox potential would be higher if the GSSG/GSH ratio were in fact greater than 80/20. A high redox potential is also implicated by the presence of ferric iron species in vacuoles [42]. Extracellular redox state can vary greatly, but in aerobic organisms and laboratory culture it is likely to be generally oxidizing compared to subcellular compartments.

The values in table 2 are not comprehensive, and should be taken as a rough guide, but even with the uncertainties, comparison with the interquartile range of *Z*_C_ of the proteins reveals some trends (figure 4*a*). The difference in both *Z*_C_ and Eh is positive going from any subcellular compartment, except for vacuoles, to the extracellular space. This pattern has an intuitive explanation: by evolutionary adjustment to optimize proteins for their environment, the inside of the cell, which is more reducing, would be expected to have more reduced proteins compared to the outside.

**Table 2:**
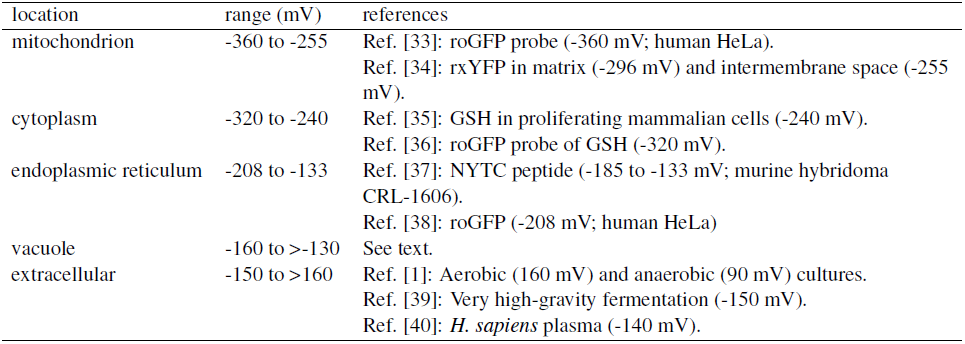
Values of Eh compiled from literature sources, for yeast cells or culture except as noted. The ranges account for variation among different cell types, experimental techniques and published values, and are used to construct figure 4.

**Figure 4:**
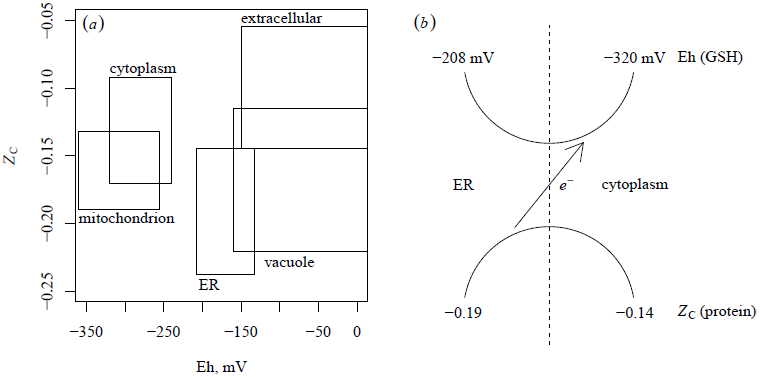
The plot (*a*) compares average oxidation state of carbon in proteins from different subcellular locations of yeast with Eh values taken from glutathione (GSH-GSSH) or other redox indicators (see table 2). The heights of the boxes indicate the interquartile ranges of *Z*_C_ values, and the widths represent the ranges of Eh listed in table 2. The scheme (*b*) invokes electron transfer to account for the contrasts in redox potential of GSH/GSSG and *Z*_C_ of proteins between ER and cytoplasm (see text).

The more surprising trend in figure 4*a* is an inverse relationship between *Z*_C_ and Eh of the cytoplasm and ER. Does this contrast have any biochemical significance? The ER is a component of the secretory pathway, which transports proteins to membranes and to outside the cell [38]. Let us conjecture that the populations of proteins in the ER and cytoplasm are connected through common metabolic intermediates – their formation and degradation are part of the recycling of biomass through endogenous metabolism [5], also implied by metabolic closure [14]. It follows that the formation of proteins of a higher *Z*_C_ in the cytoplasm entails the loss of electrons from proteins into metabolic pathways. Perhaps these pathways ultimately transfer these electrons to the formation of GSH in the cytoplasm.

This scheme is depicted in figure 4*b*. The vertical dashed line represents a physical (but not impermeable) boundary between ER and cytoplasm; the curved lines represent reactions and transport within glutathione and protein systems that cross the compartments, and the arrow represents a linkage between glutathione and protein systems, which is effectively a coupled oxidation-reduction reaction. The scheme represents a redox mass-balance interpretation of the overall stoichiometric relationships, not a mechanism applied to individual GSH or protein molecules; the complete picture of metabolic connectivity in the cell is certainly more complex. Note that this scheme refers to the relative oxidation states of carbon of the proteins, not the oxidation of protein thiol groups to form disulfides. Disulfide bond formation takes place during folding and secretion of proteins, and may also contribute to glutathione metabolism [4]. Although there is growing detail of the pathways of glutathione metabolism in the cell, including compartmentalization between ER and cytoplasm, it is not known how they connect with non-thiol systems (e.g. [40]). Integrating the oxidation-reduction requirements of protein metabolism into existing metabolic models may help to complete the balance sheet of redox interactions in the cell.

Experiments on the connections between redox conditions and protein metabolism at the subcellular level can help elucidate the possible effects of coupling of protein metabolism to the glutathione redox system. If the net transfer of high-*Z*_C_ proteins to the cytoplasm is stopped, then a decrease in GSH/GSSG redox potential in the ER relative to the cytoplasm would be expected based on the scheme shown in figure 4. This prediction is consistent with the outcome of experiments showing that puromycin-induced halting of protein synthesis causes a decrease in the redox potential monitored by roGFP in the ER [43]. A further untested implication of this hypothesis is that in ER-stress experiments the *Z*_C_ of the protein population in the ER would increase. Metabolism of proteins might also interact with redox pathways other than the oxidation and reduction of glutathione. A linkage of this type has been documented in plants, where degradation of aromatic and branched-chain amino acids was identified as a source of electrons for the mitochondrial electron transport chain [44].

### 3.4 Comparison of *Z*_C_ with standard reduction potentials of proteins

In plants, the oxidation and reduction of the iron-sulfur cluster in ferredoxin and the thiol/disulfide groups in ferredoxin-thioredoxin reductase and thioredoxin are coupled to form the ferredoxin/thioredoxin system [45] (“system” here refers to the set of interacting proteins, and is not a system in the thermodynamic sense). Ferredoxin has the lowest standard reduction potential (midpoint potential, *E*°′ at 25 °C) in this system, and its iron-sulfur cluster is reduced by light energy through photosystem I. The oxidation of ferredoxin coupled to the reduction of thioredoxin is catalysed by ferredoxin/thioredoxin reductase. Reduced thioredoxin can disseminate the redox signal via reduction of disulfide groups in other proteins, activating their enzymatic functions. Glutaredoxins are another group of thiol/disulfide proteins that also interact with disulfide bonds in proteins and, unlike thioredoxin, are reduced by glutathione [46].

The standard reduction potentials of proteins in the ferredoxin/thioredoxin system in spinach [47] and of thioredoxin and the glutaredoxin system in *E. coli* [48] are compared with *Z*_C_ in figure 5. In the ferredoxin-thioredoxin chain of spinach, there is a strong decrease in *Z*_C_ with increasing *E*°′. In the glutaredoxin system of *E. coli* and the associated proteins protein disulfide isomerase (PDI) and thiol:disulfide interchange protein (DsbA), there is a smaller decrease in *Z*__C__ with increasing *E*°′. Thioredoxin in *E. coli*, which has *Z*_C_ and *E*°′ that are similar to thioredoxin in spinach, does not follow the trend apparent in the glutaredoxin and related proteins.

**Figure 5:**
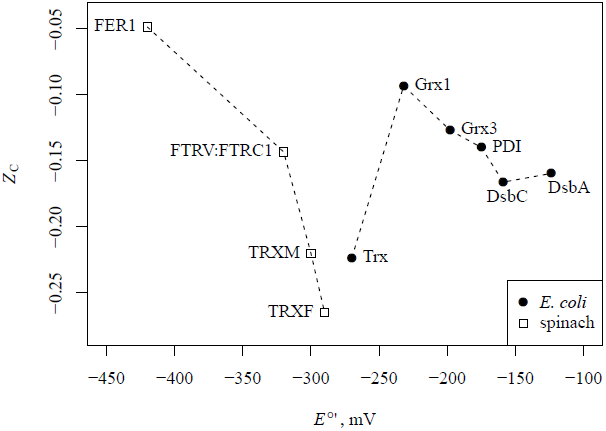
Average oxidation state of carbon in proteins compared with standard reduction potentials for ferredoxin (FER1), ferredoxin/thioredoxin reductase (FTRV:FTRC1 dimer), thioredoxin *f* (TRXF) and *m* (TRXM) from spinach [47] and thioredoxin (Trx), glutaredoxin 1 (Grx1) and 3 (Grx3), protein disulfide isomerase (PDI), and thiol:disulfide interchange protein A (DsbA) and C (DsbC) from *E. coli* [48].

From figure 5 it appears that the *Z*_C_ of proteins involved in some parts of the redox signalling networks are inversely correlated with their standard reduction potential. Do systematic changes in amino acid composition implied by *Z*_C_ impact the chemistry of the active site? The atomic environment surrounding the redox-active sites affects reduction potential [49], and changing very few residues in the vicinity of the active site can affect the function [50]. However, these or similar proximal effects may not provide a complete explanation for the trends in *Z*_C_ in figure 5, which apply to the entire protein sequences. A speculative explanation is offered here. In general, any oxidation of a protein molecule may involve loss of an electron from the active site or from a covalent bond distal to the active site. In proteins with higher *Z*_C_, the degree of oxidation of amino acids is greater, making the further loss of electrons from covalent bonds more difficult. The covalent oxidation of proteins ultimately leads to their degradation, disrupting the function of redox signalling networks. Therefore, it may be advantageous for low-*E*°′ active sites (those that have a greater potential to lose electrons on signalling timescales), to be associated with high-Z_C_ proteins (those in which the covalent bonds have lost a greater number of electrons on evolutionary timescales). The break in the pattern by thioredoxin in *E. coli*, and the scattered distribution of Z_C_ and *E*°′ of many other redox-active proteins that are not shown, presumably indicate that these hypothetical relations are applicable only to closely interacting chains of redox-active proteins and not the entire cell.

There are dual hypotheses here: first, that high-Z_C_ proteins have a lower tendency for irreversible oxidative degradation, and second, that reversible reduction potential and tendency for irreversible oxidation, which are different biochemical properties of the same molecule, are jointly tuned by evolution. One implication of the first hypothesis is that thioredoxin in spinach, which has a relatively low Z_C_, would be more easily covalently oxidized, and therefore may have a higher turnover rate than ferredoxin.

### 3.5 Phylogenetic variation in *Z*_C_ of proteins and comparison with optimal growth temperature

A comparison of Z_C_ of the combined proteins from selected microbial genomes is shown in figure 6. The sets of proteins shown on the left-hand side of figure 6 correspond to those organisms whose scientific names contain the indicated substring. In many cases, the names of the organisms reflect their environments and/or metabolic strategies. Examples of the matching genus names are *Natronobacterium*, *Haloferax*, *Rhodobacter*, *Acidovorax*, *Methylobacterium*, *Chlorobium*, *Nitrosomonas*, *Desulfovibrio*, *Geobacter*, *Methanococcus*, *Thermococcus*, *Pyrobaculum*, *Sulfolobus*. Most terms, however, match more than one genus (e.g. *<u>Pyro</u>baculum* and *<u>Pyro</u>coccus*). On the right-hand side of figure 6 are shown genera containing many groups with clinical and technological relevance; by the numbers of points it is apparent that their representation in RefSeq is greater than that of the environmental microbes.

**Figure 6:**
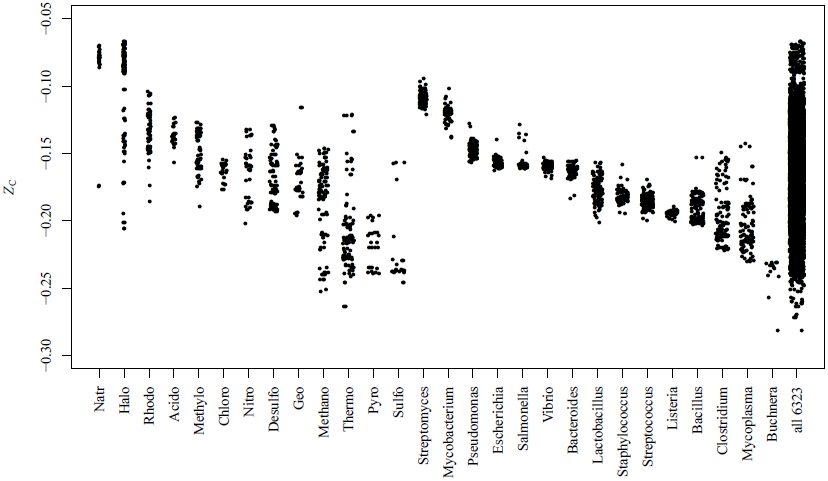
Average oxidation state of carbon in total combined proteins from sequenced microbial genomes. Sequences of microbial proteins were taken from NCBI RefSeq release 61. Only organisms with total sequenced protein length greater than 100,000 amino acids were used, leaving 6323 organisms. The group on the left-hand side is identified by substring matches in the scientific name of the organism; the terms were chosen to emphasize environmental variation. The group on the right-hand side consists of the indicated genera, emphasizing organisms of clinical and biotechnological relevance. The final category represents total proteins from all microbial genomes meeting the minimal size requirement, many of which are not shown in the other categories.

A general trend toward lower *Z*_C_ in proteins in organisms from hot environments (e.g. represented by Thermo and Pyro in the names) is apparent. Organisms with the highest *Z*_C_ inhabit saline evaporative waters (Natr, Halo), while other aquatic organisms (e.g. Rhodo) have less highly oxidized proteins. Within a given genus (right-hand side of plot), the clusters of *Z*_C_ tend to be tighter, reflecting conserved compositional trends. *Streptomyces*, common in soils, has the highest *Z*_C_ of the genera shown here. *Buchnera* is notable because its proteins are highly reduced, including one example (*Buchnera aphidicola* BCc) with the lowest *Z*_C_ of proteins within the entire dataset. *B. aphidicola* BCc is the primary endosymbiont of the cedar aphid *(Cinara cedri)* and, at the time of sequencing, had one of the smallest known bacterial genomes [51]. Being relatively closely related [52], *Mycoplasma* and *Clostridium* also have relatively low *Z*_C_. *Mycoplasma are* known for their small genomes and dependence on metabolic products of the host; the low *Z*_C_ may be a constraint imposed by growth in reducing intracellular or intraorganismal environments.

We now turn to a comparison of homologues of a specific protein. Rubisco is an essential enzyme for carbon fixation. Sequence comparison of homologues (related sequences that appear in different organisms) has provided the basis for many phylogenetic studies [53]. Major divergent forms of the enzyme are Forms I and II, found in aerobic organisms, and Form III and “Rubisco-like proteins”, found in anaerobic organisms [54]. The organisms listed in table 3 have in common the occurrence of Rubisco in their genome. Forms I, II or III were included in this comparison, but Rubisco-like proteins were excluded; however, some that were tested were found to have considerably lower *Z*_C_. The selection of organisms was made in order to represent a variety of optimal growth temperatures (*T*_opt_) as reported in the studies cited in the table [55–81].

**Table 3:** Names of species, optimal growth temperatures (*T*_opt_) and UniProt [26] accession numbers (IDs) for the large subunit of ribulose bisphosphate carboxylase (Rubisco). Literature references for *T*_opt_ are indicated in brackets. Abbreviations: A – Archaea; B – Bacteria; E – Eukaryota. The numbers are used to identify the points in figure 7 (duplicated numbers occur in different temperature ranges).

| number | domain | species | $T_{\text{opt}}$, °C | reference | ID |
| --- | --- | --- | --- | --- | --- |
| 1 | E | *Phaeocystis antarctica* | 0–6 | [55] | G9FID8 |
| 2 | B | *Octadecabacter antarcticus* 307 | 4–10 | [56] | M9R7V1 |
| 3 | A | *Methanobolus psychrophilus* | 18 | [57] | K4MAK9 |
| 4 | A | *Methanococcoides burtonii* | 23 | [58] | Q12TQ0 |
| 5 | B | *Bradyrhizobium japonicum* | 25–30 | [59] | Q9ZI34 |
| 6 | B | *Thiobacillus ferrooxidans* | 28–30 | [60] | P0C916 |
| 7 | E | *Zea mays* | 30 | [61] | P00874 |
| 8 | B | *Mariprofundus ferrooxydans* | 30 | [62] | Q0EX22 |
| 9 | B | *Desulfovibrio hydrothermalis* | 35 | [63] | L0RHZ1 |
| 1 | A | *Methanosarcina acetivorans* | 35–40 | [64] | Q8THG2 |
| 2 | B | *Acidithiobacillus caldus* | 45 | [65] | F9ZLP0 |
| 3 | E | *Cyanidium caldarium* | 45 | [66] | P37393 |
| 4 | B | *Sulfobacillus acidophilus* | 45–50 | [67] | P72383 |
| 5 | B | *Pseudomonas hydrognotermophila* | 52 | [68] | Q51856 |
| 6 | B | *Synechococcus* sp. (strain JA-2-3B'a(2-13)) | 50–55 | [69] | Q2JIP3 |
| 7 | A | *Methanosaeta thermophila* | 55–60 | [70] | A0B9K9 |
| 8 | B | *Thermosynechococcus elongatus* | 57 | [71] | Q8DIS5 |
| 9 | B | *Clostridium clariflavum* | 60 | [72] | G8LZL2 |
| 1 | B | *Bacillus acidocaldarius* | 65 | [73] | F8IID7 |
| 2 | B | *Thermotoga lettingae* | 65 | [74] | A8F7V4 |
| 3 | B | *Thermomicrobium roseum* | 70 | [75] | B9KXE5 |
| 4 | A | *Archaeoglobus fulgidus* | 76 | [76] | O28635 |
| 5 | A | *Methanocaldococcus jannaschii* | 85 | [77] | Q58632 |
| 6 | A | *Thermophilum pendens* | 85–90 | [78] | A1RZJ5 |
| 7 | A | *Staphylothermus marinus* | 85–92 | [79] | A3DND9 |
| 8 | A | *Pyrococcus horikoshii* | 98 | [80] | O58677 |
| 9 | A | *Pyrococcus furiosus* | 100 | [81] | Q8U1P9 |

A comparison between *T*_opt_ and *Z*_C_ of Rubisco is presented in figure 7. The *Z*_C_ of Rubisco are somewhat higher than the bulk protein content of the organisms; compare for example the values for *Pyrococcus horikoshii* and *P. furiosus* (the highest-temperature points labelled 8 and 9 in figure 7*a*) with the range of values for “Pyro” in figure 6. At lower temperatures (0 to 50 °C), the differences between domains of life are most apparent; Rubiscos of the Bacteria in this sample set are more oxidized than those of Archaea and Eukaryota. There is a tendency for the Rubiscos of the Archaea to have lower *Z*_C_; this appears to be characteristic of anaerobic methanogenesis and Form III Rubiscos. An interesting exception is the high-*Z*_C_ Rubisco of *Methanosaeta thermophila*; this organism grows on acetate to produce both CH_4_ and CO_2_ [70]. The major pattern that emerges is that higher temperatures are associated with a lower average oxidation state of carbon in proteins. As outlined below, a decrease in oxidation state of carbon in the covalent structure of the proteins confers energetic savings in hot, reducing environments.

**Figure 7:**
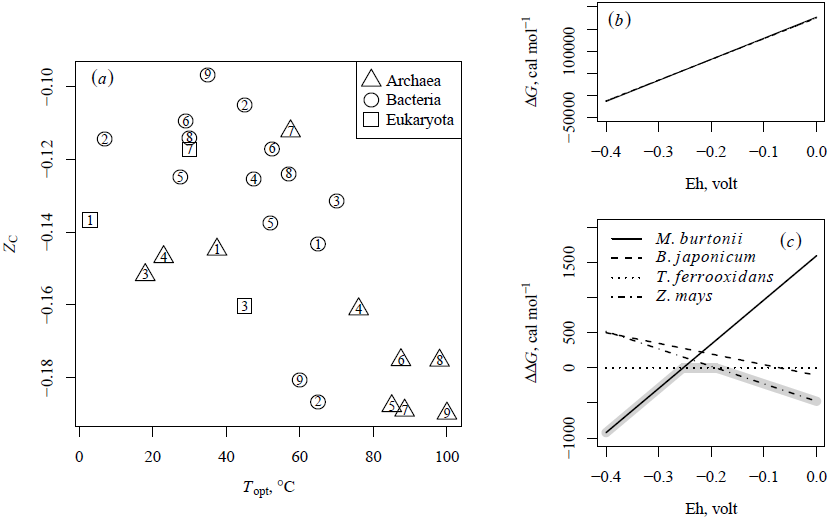
(*a*) Average oxidation state of carbon in Rubisco compared with optimal growth temperature (*T*_opt_) of organisms. Numbers are used to identify the organisms (see table 3). (*b*-*c*) Gibbs energies of formation reactions, per residue, of selected Rubisco from organisms in the 23–30 °C range of optimal growth temperature (points labelled 4–7). (*b*) Total Gibbs energies of individual reactions as a function of Eh and (*c*) difference between the reaction for *T. ferrooxidans* and the others are shown. The grey highlight indicates the protein with the lowest Δ*G* along the range of Eh values.

### 3.6 Beyond *Z*_C_: energetics of protein formation as a function of environmental redox potential

To a first approximation, energetic considerations predict that more reducing conditions tend to favour formation of proteins with relatively lower *Z*_C_, and vice versa. To assess the directionality and magnitude of chemical forces on the evolutionary transformations of proteins, the energetics of reactions can be calculated using thermodynamic models. Because the timescales of evolution are much longer than transformations of biomolecules during metabolism, a discussion of the assumptions underlying the application of thermodynamic theory to biochemical evolution is warranted.

The calculation of equilibrium provides a quantitative description of the state of a system in an energy minimum. The assumption of equilibrium is the foundation for many models of inorganic processes in geochemistry. Although the metabolic formation of proteins and other biochemical constituents proceeds in a non-equilibrium manner, using the equilibrium state as a frame of reference makes it possible to compare the energetics of living systems quantitatively (“how far from equilibrium”).

In energetic terms, adaptation can be defined as a problem of optimal efficiency, with trade-offs between energy utilization and power [82, p. 140]. Overall minimization of biomass synthesis costs can be expected from the energy utilization standpoint. The links between energetics and evolutionary outcome implicitly depend on the proposal that greater fitness is associated with mutations that lower the synthesis costs of proteins for a given function (e.g. [83]).

It has been argued previously that the propensity for some evolutionary changes can be modelled using the related concepts of equilibration, energy minimization, and maximum entropy. In an evolutionary context, these concepts have often been defined by analogy to their definitions in thermodynamics [82, 84]. The current discussion instead considers the possibility of direct application of chemical thermodynamics (the geochemical approach) to formulate a quantitative description of patterns of protein composition. The major assumption used in the following discussion is that energetic demands of protein formation depend not only on the composition of the protein, but also the environmental conditions. Therefore, it is expected that environmental adaptation has left an imprint on chemical compositions of phylogenetically distinct proteins.

An example calculation is carried out here for selected Rubiscos from organisms with optimal temperatures in the range of 23–30 °C (table 3, numbers 4–7). The basis species (representing inorganic starting materials) and their chemical activities used for this example are CO_2_ (10^‒3^), H_2_O (1), NH_3_ (10^−4^), H_2_S (10^‒7^) and H^+^ (10^‒7^, i.e. pH = 7). The activity of the electron, representing the effects of the redox variable (Eh) through the Nernst equation, is left to vary. The Gibbs energies of the reactions to form the proteins were calculated as described previously [9, 19] and are plotted in figure 7*b*. In figure 7*b* it can be seen that the Gibbs energies of the reactions to form the proteins (Δ*G*), normalized by protein length, steadily increase (become less favourable) with increasing Eh, but the differences between the proteins can not be discerned easily. In figure 7*c*, the values of Δ*G* are shown relative to the reaction for *T. ferrooxidans*, giving a difference in Gibbs energy of formation (ΔΔ*G*) that can be used to assess the relative energetics of the reactions. Lower energies indicate the most stable protein (in the sense of chemical formation, not structural conformation), and these relative energies depend on the chemical conditions of the environment, measured in part by the oxidation-reduction potential.

By using the Gibbs energy calculations such as shown in figure 7*c*, one can make an assessment of which protein in a given redox environment demands lower energy for overall synthesis (i.e. is more stable) than other possibilities. Where two lines cross in figure 7*c*, the energies to form the two proteins are equal, representing a metastable (partial) equilibrium. In metastable equilibrium, the coexistence of proteins with equal energies of formation corresponds to a local energy minimum. This interpretation does not preclude the non-equilibrium character of the overall biosynthetic process, because the energies of formation remain non-zero (figure 7*b*).

The three least costly proteins, going from low to high Eh, are those from *M. burtonii, T. ferrooxidans*, and *Z. mays*. Methanogenic archaea such as *M. burtonii* [58] inhabit reducing environments, where Eh values as low as at least −400 mV have been documented [2]. In contrast, the metal-leaching activity of *T. ferrooxidans* is associated with an increase in oxidation-reduction potential (ORP, which carries the same units as Eh) [85] and, compared to *M. burtonii*, its Rubisco is more oxidized and is consequently more stable at higher Eh. The correspondence between redox conditions and lower Δ*G* of formation of the proteins provides thermodynamic evidence for environmental adaptation of the proteins.

The protein from *T. ferrooxidans*, which has the highest *Z*_C_ of the four considered, does not have the lowest value of ΔΔ*G* in the most oxidizing (highest Eh) conditions. Instead, the Rubisco from *Z. mays*, even though it has lower *Z*_C_, is calculated to relatively more stable at the highest Eh values considered. This result shows that comparing values of *Z*_C_ provides only an approximation of the dependence of the relative energetics of protein formation on redox changes; chemical thermodynamic models integrate more information about reaction stoichiometry and the effects of multiple environmental variables.

Previously, comparison of Rubisco from hot-spring organisms adapted to higher temperatures revealed an increase in frequency hydrophobic amino acids, interpreted as increasing the conformational stability of proteins at high temperature [86]. However, another basis for interpretations of thermophilic adaptation depends on the relative energetic costs of synthesis of amino acids [87]. The finding made in this study is that the high-temperature Rubiscos exhibit a shift toward lower *Z*_C_ (figure 7*a*). Higher temperatures are often associated with more reducing environments. For example, compared to seawater, activities of dissolved hydrogen in hydrothermal fluids are higher, and mixed hydrothermal-seawater fluids have a reducing potential that favours formation of relatively reduced amino acids [8]. Redox potential is a major variable affecting the energetics of protein formation at different temperatures; therefore, adaptation to minimize biosynthetic costs in high-temperature environments is likely to have more than just a thermophilic (temperature dependent) aspect.

Although the stoichiometric comparisons are distinct from sequence-based phylogenetic analyses which are used to test for positive selection, it is conceivable that *Z*_C_ could in the future be incorporated into tree-based models of evolution that take account of physicochemical properties of amino acids in proteins [88]. However, as noted above, thermodynamic comparisons have greater power than compositional comparisons such as *Z*_C_. The computation of metastable equilibrium takes account simultaneously of temperature, redox potential (expressed as Eh, activity of hydrogen, or oxygen fugacity) and other variables [19]. In another study, analysis of metagenomic and geochemical data led to a predicted metastable succession of proteins (with generally increasing *Z*_C_) that could be aligned with a gradient of increasing oxidation potential and decreasing temperature in a flowing hot spring [10]. When grouped by taxonomic similarity, the *Z*_C_ in the hot-spring proteins, while becoming lower at high temperature, also spread over a broader range, leading to tighter constraints on the redox conditions suitable for metastable coexistence of the organisms [11].

Not only differences in the oxidation states of present-day environments, but also the oxygenation of Earth’s atmosphere and oceans through geological time could have profound impacts on the energetics of biomass synthesis [7]. Adaptation to reduce these costs likely would lead to divergences in *Z*_C_ that are apparent across different taxa, while closer phylogenetic relationships should confer a similarity in *Z*_C_. In common with the Rubiscos, comparison of the total proteins of microbes reveals a tendency for proteins in organisms associated with hot, as well as sulfidic and methanogenic environments, to be more reduced (figure 6).

## 4 Conclusions

Proteins are products of metabolism; their synthesis and degradation are part of the network of chemical reactions that sustains the living cell. Comparisons of the average oxidation state of carbon in proteins have provided a starting point for visualizing the compositional diversity of proteins in relation to redox chemistry in subcellular compartments and external environments. The large differences in *Z*_C_ of proteins in locations such as the ER and cytoplasm likely have consequences for the dynamics of oxidation-reduction reactions involving glutathione and other metabolites. Further insight may be gained by including the formation and degradation of proteins in kinetic and stoichiometric models of metabolic networks. Extension of these concepts to other phenomena entailing changes in both redox potential and protein expression, such as stress response to oxidizing agents and the cell cycle, can be envisioned. The deepest significance of the observed patterns lies in their emergence over evolutionary timescales. The inverse trend relating *Z*_C_ with standard reduction potentials in chains of redox-active proteins is a case where the chemical composition of the proteins may be tuned with the electron-transfer chemistry of the active sites. Compositional divergences among proteins are also apparent in phylogenetic comparisons, and here it is reasonable to conclude that correlations between oxidation state of carbon in proteins and the redox potential of the environment indicate some degree of energy savings conferred by evolution. The natural history of protein evolution is a result of processes that are both unpredictable (mutation events) and, to some extent, deterministic (selection for fitness in a given environment). By describing protein molecules in terms of chemical composition and energetics it will be possible to identify some of the forces that help to shape the occurrences of proteins in different cells and environments.

## 5 Supporting information

The supporting information is provided in a ZIP archive containing the following code and data files:

**prep.R** This file contains code used to prepare the data files for easier handling by the plotting functions.
**plot.R** This file contains code used to make the figures appearing in this paper. The functions, in the order of the figures (1–7), are amino(), human(), yeast(), potential(), midpoint(), phylo(), and rubisco(). The code is written in R [18] and depends on version 1.0.2 of the CHNOSZ package [19], available from the Comprehensive R Archive network (http://cran.r-project.org).
**data/SGD_associations.csv** For yeast genes, this table lists the accessions, SGDID, and the association to cellular components in the Gene Ontology, derived from gene_association.sgd.gz, pro-tein_properties.tab and go_terms.tab downloaded from http://www.yeastgenome.org on 2013-08-24. All gene associations with the NOT qualifier were removed, as were those without a matching entry in protein_properties.tab (e.g. RNA-coding genes).
**data/ZC_HUMAN.csv,ZC_membrane.csv** Compilations of the values of *Z*_C_ for human proteins and human membrane proteins. Values in ZC_HUMAN.csv ware calculated from protein sequences in HUMAN.fasta.gz, downloaded from ftp://ftp.uniprot.org/pub/databases/uniprot/current_release/knowledgebase/proteomes/HUMAN.fasta.gz on 2013-08-24 (file dated 2013-07-24). Values in ZC_membrane.csv were calculated from protein sequences in all *.fa files in Additional File 2 of Almén et al., 2009 [27].
**data/codons.csv** In the first column, the three-letter abbreviations for each of the RNA codons; in the second column, the names of the corresponding amino acids.
**data/midpoint.csv** List of protein names, UniProt IDs and standard midpoint reduction potentials used to make figure 5. Start and stop positions, taken from UniProt, identify the protein chain excluding initiator methionines or signal peptides.
**data/protein_refseq.csv** Amino acid compositions of total proteins in 6758 microbial genomes from RefSeq release 61, dated 2013-09-09. The gene identifier (gi) numbers of the sequences were assigned taxonomic IDs (taxids) using the RefSeq release catalogue. The amino acid compositions of the total proteins were calculated by averaging the compositions of all proteins for each taxid. The “organism” column contains the taxid used in NCBI databases, the “ref” column contains the names of the RefSeq files from which the amino acid sequences were taken (with start and end positions in parentheses) followed by the scientific name of the organisms in brackets, and the “abbrv” column contains the number of amino acids for that organism. Scientific names for the taxids at the species level were found using the names.dmp and nodes.dmp files downloaded from ftp://ftp.ncbi.nih.gov/pub/taxonomy/taxdump.tar.gz on 2013-09-18.
**data/rubisco.csv** UniProt IDs for Rubisco and optimal growth temperatures of organisms (see table 3).
**cell/*.png** PNG images for each of the cellular components used to make figure 3.
**fasta/midpoint/*.fasta** FASTA sequence files for proteins shown in figure 5.
**fasta/rubisco/*.fasta** FASTA sequence files for each Rubisco identified in table 3.

## Acknowledgements

Thanks to Svenja Tulipani and Katy Evans for their comments on an earlier version of the manuscript.

